# Mechanoregulatory role of TRPV4 in prenatal skeletal development

**DOI:** 10.1101/2022.06.23.497086

**Authors:** Nidal Khatib, James Monsen, Saima Ahmed, Yuming Huang, David A. Hoey, Niamh C. Nowlan

## Abstract

Biophysical cues are essential for guiding skeletal development, but the mechanisms underlying the physical regulation of cartilage and bone formation are unknown. TRPV4 is a cell membrane ion channel responsible for transducing mechanical stimuli as a means of regulating skeletal cell homeostatic processes. Dysregulation of TRPV4 is associated with several skeletal developmental pathologies, indicating its involvement in cartilage and bone development, potentially in a mechanosensing capacity. In this study, we test the hypothesis that mechanically mediated prenatal skeletogenesis depends on TRPV4 activity. We first validate a novel model where we establish that dynamically loading embryonic mouse hindlimb explants cultured *ex vivo* promotes joint cartilage growth and morphogenesis, but not diaphyseal mineralization. We next reveal that TRPV4 protein expression is mechanically regulated and spatially localized to patterns of high biophysical stimuli in the femoral condyles of cultured limbs. Finally, we demonstrate that TRPV4 activity is crucial for the mechanical regulation of joint cartilage growth and shape, mediated via the control of cell proliferation and matrix biosynthesis, indicating a mechanism by which mechanical loading could direct morphogenesis during joint formation. We conclude that the regulatory pathways initiated by TRPV4 mechanotransduction are essential for the for the cartilage response to physical stimuli during skeletal development. Therefore, TRPV4 may be a valuable target for the development of therapeutic skeletal disease modifying drugs and developmentally-inspired tissue engineering strategies for skeletal repair.

## Introduction

Skeletogenesis begins in early embryonic development when the entire skeleton forms first as a cartilaginous template (Hall & Miyake, 2000; Karsenty, 2003). The cartilage anlagen are gradually mineralised over prenatal and postnatal development to form the mature skeleton (Kronenberg, 2003). The ultimate size, shape and organisation of skeletal tissues depends on many cellular processes, including matrix production, cell division, hypertrophy, intercalation, as well as changes in cell size and orientation (Brunt et al., 2016; Decker et al., 2017; Godivier et al.; Shwartz et al., 2012; Zhang et al., 2020). The spatial and temporal orchestration of such a complex process depends on an interplay between genetic and epigenetic mechanisms influenced by environmental factors, notably mechanical signals (Nowlan, Sharpe, et al., 2010; Pollard et al., 2014). Skeletogenesis in later development is influenced by fetal movements such as kicking, which induce stresses and strains on the developing skeleton (Verbruggen et al., 2018). Reduced or absent fetal movement is associated with skeletal abnormalities such as hip dysplasia, joint contractures (fixed joints) and slender, hypo-mineralised bones (Aronsson et al., 1994; Bacino et al., 1993; Rodriguez et al., 1988). Animal models of fetal immobilization or reduced muscle display skeletal abnormalities akin to that found in akinesia syndromes, including shorter limbs with reduced bone, failed joint cavitation and joint shape malformation (Bridglal et al., 2021; Brunt et al., 2016; Kahn et al., 2009; Nowlan, Bourdon, et al., 2010; Nowlan et al., 2014; Roddy et al., 2011; Sotiriou et al., 2019). While substantial progress has been made mapping the genetic molecular control of skeletal development (Karsenty & Wagner, 2002; McQueen & Towers, 2020), we still have a poor understanding of the mechanisms underlying the influence of mechanical factors. Identifying key mechanoregulatory mechanisms of skeletogenesis would be valuable for advancing our understanding of biophysical developmental control and motivate developmentally inspired mechanotherapeutic or skeletal tissue regeneration strategies.

Skeletal cells such as chondrocytes and bone cells sense and respond to mechanical stimuli as a means of regulating skeletal tissue growth, homeostasis and repair (Wang & Thampatty, 2006). Mechanical loading of skeletal tissues exposes cells to multiple modes of stress, such as compression, tension, hydrostatic pressure, osmotic pressure and fluid shear. In a process called mechanotransduction, biophysical cues are detected by specialized mechanosensors that convert them into biochemical signals, consequently regulating cell behaviour (Dieterle et al., 2021; Lewis et al., 2017; Liedtke & Kim, 2005). Cell membrane mechanosensors detect physical changes at the cell membrane, such as integrins which sense cell-matrix interactions through focal adhesions with the substrate (Wang et al., 1993), cadherins which transduce cell-cell interactions (Wang et al., 2009), and mechanosensitive ion channels which activate in response to physical changes in the membrane such as tension (Syeda et al., 2016), hypotonicity (Liedtke & Friedman, 2003), deflection of cell-matrix focal adhesions (Servin-Vences et al., 2017) and deflection of the primary cilia (Corrigan et al., 2018). While integrins, cadherins and ion channels have been shown to regulate homeostatic cell activities such as proliferation, survival, maturation and matrix deposition in skeletal cells (McMahon et al., 2008; Mobasheri et al., 2002; Mouw et al., 2007), their involvement in mechanically regulated processes of cartilage and bone development remains unclear.

The involvement of mechanosensitive ion channels in skeletal development has been highlighted by the effects of hereditary functional mutations (channelopathies) of Transient receptor potential vallinoid 4 (TRPV4). TRPV4 is a polymodal Ca^2+^ permeable non-selective ion channel (Liedtke, 2007), mutations in which leads to a phenotypically diverse range of severe skeletal conditions including lethal metatropic dysplasia, spondylometaphyseal dysoplasia (dwarfirm), and autosomal dominant brachyolmia (Camacho et al., 2010; Nilius & Voets, 2013; Nishimura et al., 2012). First discovered in 2000 (Liedtke et al., 2000; Strotmann et al., 2000), TRPV4 was initially found to be responsible for transducing osmotic signals (Liedtke & Friedman, 2003), but has since been implicated in the cell biosynthetic response to compressive loading (O’Conor et al., 2014), hydrostatic pressure (Savadipour et al., 2022) and oscillatory fluid shear (Corrigan et al., 2018). Previous work has found a critical role for TRPV4 in cartilage and bone cell function *in vitro*. Chemical activation of the ion channel using specific channel agonists stimulates upregulation of chondrogenic gene expression markers (*ACAN, Col2a1)* and the production of cartilaginous matrix proteins glycosaminoglycans and collagen (Corrigan et al., 2018; O’Conor et al., 2014; Willard et al., 2021). Upregulated TRPV4 activity has also been associated with chondrogenic differentiation of iPSCs, MSCs and iPSC-derived chondroprogenitors (Adkar et al., 2019; Huynh et al., 2019; O’Conor et al., 2014; Willard et al., 2021), indicating a role of TRPV4 in chondrogenesis. Interestingly, *Trpv4*^*-/-*^ knockout mice develop spontaneous osteoarthritis at a younger age than wild type controls (Clark et al., 2010), while cartilage-specific *Trpv4*^*-/-*^ mice experience reduced severity of aging-associated osteoarthritis than wild type controls (O’Conor et al., 2016). In bone, TRPV4-mediated signalling has been shown to regulate osteocyte differentiation (Masuyama et al., 2008), while *Trpv4*^*-/-*^ knockout mice exhibit reduced bone loss and reduced osteoclast function in an unloading-induced bone loss model (Mizoguchi et al., 2008), suggesting a key involvement of TRPV4 in regulating bone adaptation to loading. While previous work has revealed an involvement of TRPV4 in both skeletal developmental diseases and mature skeletal cell activity *in vitro*, the specific function of the ion channel has not yet been characterised in developing skeletal tissues.

Major advances in TRPV4 agonist and antagonist discovery in recent years and successful evaluation of the first selective TRPV4 antagonist in clinical trials (Goyal et al., 2019; Lawhorn et al., 2020) provides a promising view of the development of human therapies. Furthermore, TRPV4 has been recognized as a potentially valuable therapeutic target for joint diseases (McNulty et al., 2015; Nilius & Szallasi, 2014), therefore, a better understanding of how TRPV4 functions in skeletal tissue development may be critical for informing therapies targeting TRPV4 mutation-associated disease or skeletal regeneration in the near future. In this study, we test the hypothesis that mechanically regulated cartilage formation and mineralization of developing hindlimb skeletal tissues are mediated by TRPV4 mechanotransduction. We first quantify the effects of mechanical loading on cartilage growth, morphogenesis and diaphyseal mineralization of embryonic murine hindlimb explants using an *in vitro* mechanostimulation bioreactor system. Next, we determine if mechanoregulation of skeletal development is dependent on TRPV4 activity. Finally, we elucidate the involvement of TRPV4 in the physiological link between mechanical loading and cartilage growth, cell proliferation and matrix biosynthesis during skeletal development.

## Results

### Dynamic stimulation of mouse embryo limb explants promotes cartilage growth and morphogenesis, but not diaphyseal mineralization

The first step of this research established the effects of dynamic mechanical loading on cartilage growth, shape and mineralization of hindlimb explants from mouse embryos. We previously developed a bioreactor system to apply dynamic mechanical loading regimes to embryonic chick limb explants *ex vivo*, establishing a positive effect of loading on joint morphogenesis (Chandaria et al., 2016; Khatib et al., 2021) and diaphyseal mineralization (Khatib et al., 2021). This model permits rigorous control over the mechanical stimuli applied to developing skeletal tissues, a major challenge in *in ovo* animal model studies (Nowlan, Sharpe, et al., 2010). In this study, we utilised mouse embryo hindlimbs in our mechanostimulation bioreactor system which permits investigation of the distinct mammalian processes of bone formation from avian development, and opens up avenues for culturing limbs from transgenic mice. To determine the effect of dynamic culture on skeletal development *in vitro*, embryonic day 15.5 mouse hindlimbs were cultured for 6 days, with one limb in dynamic culture conditions and the contralateral limb in static conditions (Fig 1A, 1B and supplementary video). Limbs were stained with Alcian blue (cartilage) or alizarin red (mineral), imaged in 3D using optical projection tomography (OPT), and then segmented to generate 3D models (Quintana & Sharpe, 2011). Rudiment length and joint shape parameters were measured from the models (Fig 1C).

**Figure 1:**
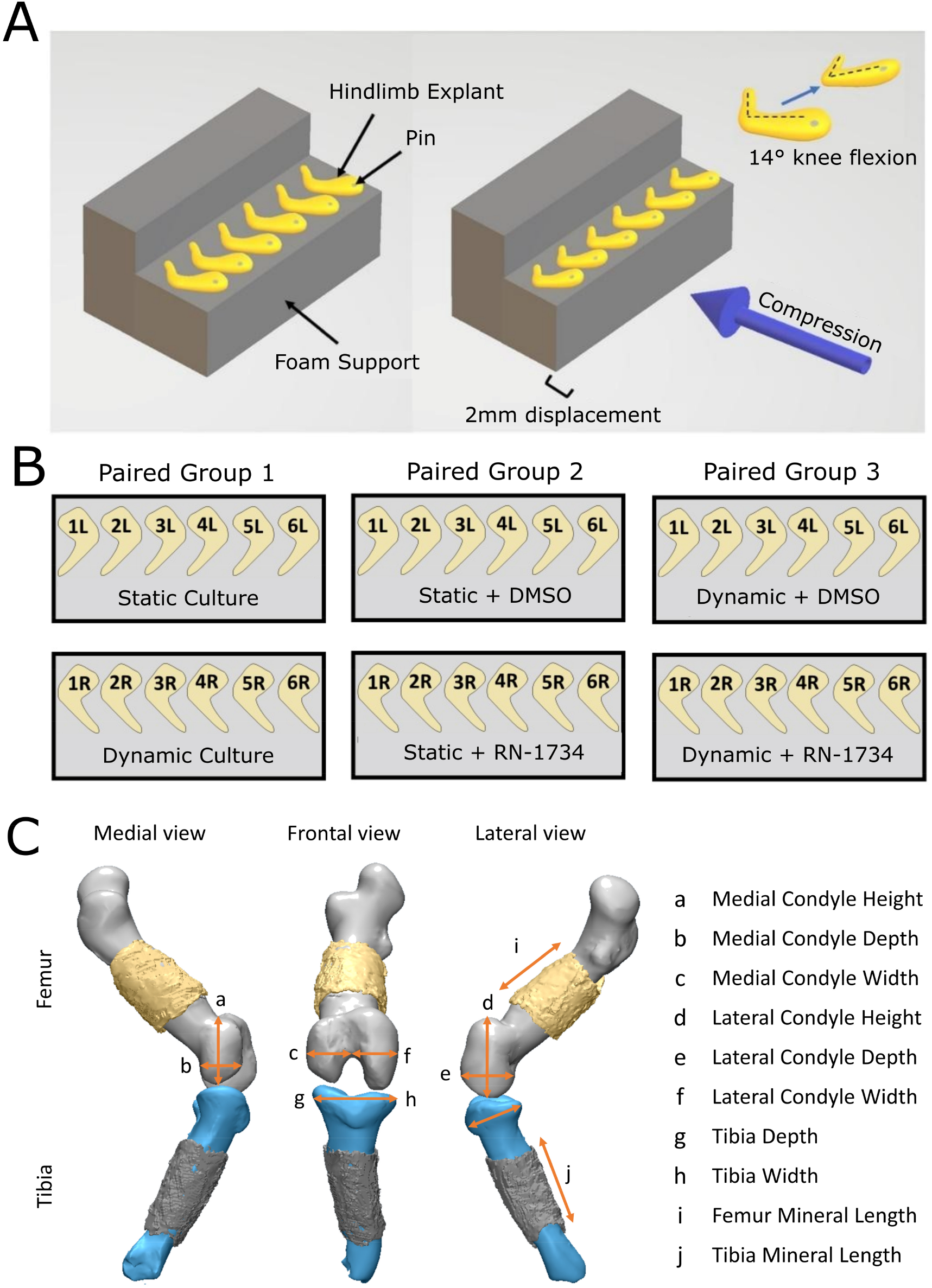
Experimental set up for mechanical stimulation of mouse hindlimbs, comparison groups, and visualization of measurements for quantitative analysis. (A) Six to eight limbs pinned to foam supports were placed within bioreactor chambers for dynamic culture or petri dishes for static culture. Dynamic cultured limbs were exposed to cyclic flexion-extension movements of approximately 14 ° (±2 °) at 0.67 Hz, applied by compressive displacement of the foam supports. (B) Within each comparison group, contralateral limbs from the same embryo served as paired samples for three comparison groups. (C) Eight cartilage joint features and two mineral length measurements were measured from 3D cartilage and mineral models generated from OPT data.

Mechanical loading induced by dynamic culture significantly increased growth of six out of eight knee joint shape features compared to static controls (*p*<0.05, n=8), namely the width, height and depth of the lateral condyle, tibial depth and width, and medial condyle depth (Fig 2A and S1.1). Medial condyle height and width were not significantly increased by dynamic loading, but trends of increased growth of these parameters with loading were evident (Fig 2A and S1.1). Visual analysis of the shapes outlines revealed that the most prominent effects of dynamic culture on joint shape were increased definition of the posterior curl of the lateral condyle (Fig 2C, *), and medio-lateral expansion of the lateral condyle (Fig. 2C, †). From these data we conclude that growth and morphogenesis of the joint cartilage of cultured explants are influenced by mechanical loading in our culture system.

**Figure 2:**
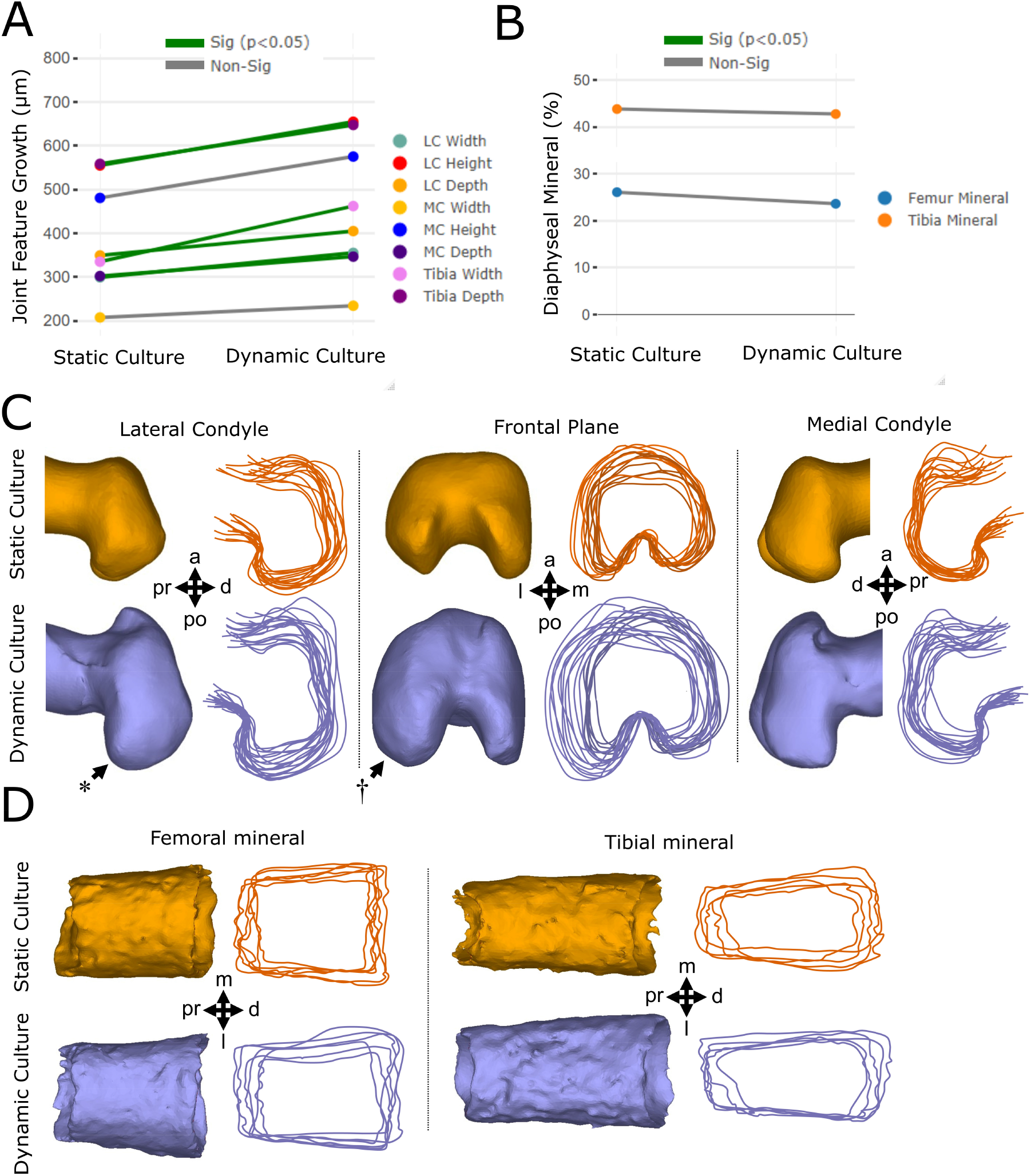
Dynamic loading of cultured mouse embryo hindlimbs promotes joint cartilage feature growth and shape, but not the extent of mineralization or bone collar shape. (A) Average paired differences in growth for joint cartilage features (each line represents one of eight features) in static versus dynamic culture (n=8 limbs per group). Green lines; significant (*p<*0.05) feature differences. Gray lines; non-significant feature differences. (B) Average extents of femur (n=6) and tibia mineralization (n=6) in static and dynamic cultured paired limbs. (C) Representative samples and joint shape contours for distal femora cultured in static and dynamic conditions. Arrows indicate regions of increased growth and shape development. * represents more prominent posterior curl observed in dynamically loaded limbs. † represents larger lateral expansion found in the lateral condyle of dynamically loaded limbs. (D) Representative samples and outlines of femoral and tibial bone collars from limbs cultured in static and dynamic conditions. Pr; proximal, d; distal, a; anterior, po; posterior, m; medial, l; lateral.

In contrast to reported increases in the diaphyseal mineralization of chick limb explants in response to mechanical stimuli (Henstock et al., 2013; Khatib et al., 2021), dynamic loading of cultured murine limbs had no significant effects on diaphyseal femoral or tibial mineral length (Fig 2B and S2). Furthermore, no visible differences were observed in bone collar shape between dynamically and statically cultured mouse femora or tibiae (Fig 2D). Together, our findings indicate that dynamic loading of *ex vivo* mouse embryo limbs stimulates cartilage growth and morphogenesis, but not mineralization. The differences found in the mineralization response to loading across the mouse and chick model is likely due to the reliance of endochondral ossification on blood vessel invasion in the mammalian limb (Kronenberg, 2003; Nowlan et al., 2007). In contrast to mammalian osteogenesis, avian long bones do not require vascularization of the primary cartilage prior to mineralization (Nowlan et al., 2007). Since the vasculature in our hindlimb explants becomes non-functional after dissection, it is possible that osteogenic progenitors do not reach the ossification sites in our murine cultured hindlimbs. In summary, we have validated a powerful new model of *ex vivo* murine limb development which captures key stages of mechanically driven cartilage growth and joint morphogenesis.

### TRPV4 protein expression is mechanically regulated in developing murine cartilage

TRPV4 protein expression has been identified in most adult skeletal cell types (Masuyama et al., 2008), but the variance and patterning of its expression in developing skeletal tissues has not previously been investigated. To characterise the spatial protein expression of TRPV4 in prenatal skeletal tissues and determine how it is affected by *ex vivo* loading, we compared the localization of TRPV4 protein concentrations between statically and dynamically cultured hindlimb explants. Cultured femora were sectioned and labelled for TRPV4 at the protein level using immunofluorescence staining. TRPV4 protein was found to be co-localization to the cell membrane in murine chondrocytes (Fig S3). Three regions of the femur were imaged and quantitatively analysed, namely the medial condyle, lateral condyle and growth plate. Individual cell TRPV4 intensities were measured across all chondrocytes captured within cropped areas of these three regions.

In statically cultured limbs, TRPV4 localisation was evident in the hypertrophic cartilage of the growth plate, but not prominent in the metaphyseal and joint cartilage (Fig 3A). Dynamic loading led to a dramatic increase in the prominence of TRPV4 protein expression in the medial and lateral condyles compared to limbs cultured in static conditions, while expression in the growth plate was not affected by loading when comparing across all samples (Fig 3A and S4). TRPV4 protein expression was spatially varied throughout the dynamically loaded condyles, with increased expression in the anterior aspect of the medial condyle and in the anterior and proximal aspects of the lateral condyle. Quantitative data on TRPV4 intensity revealed a significant increase in intensity for the medial and lateral condyles due to dynamic loading, but no change in intensity at the growth plate (Fig 3B). These data demonstrate that in *ex vivo* cultured embryonic limb explants, TRPV4 protein concentration is mechanically upregulated in joint epiphyseal chondrocytes, while regulation of TRPV4 in hypertrophic chondrocytes is independent of extrinsic loading.

**Figure 3:**
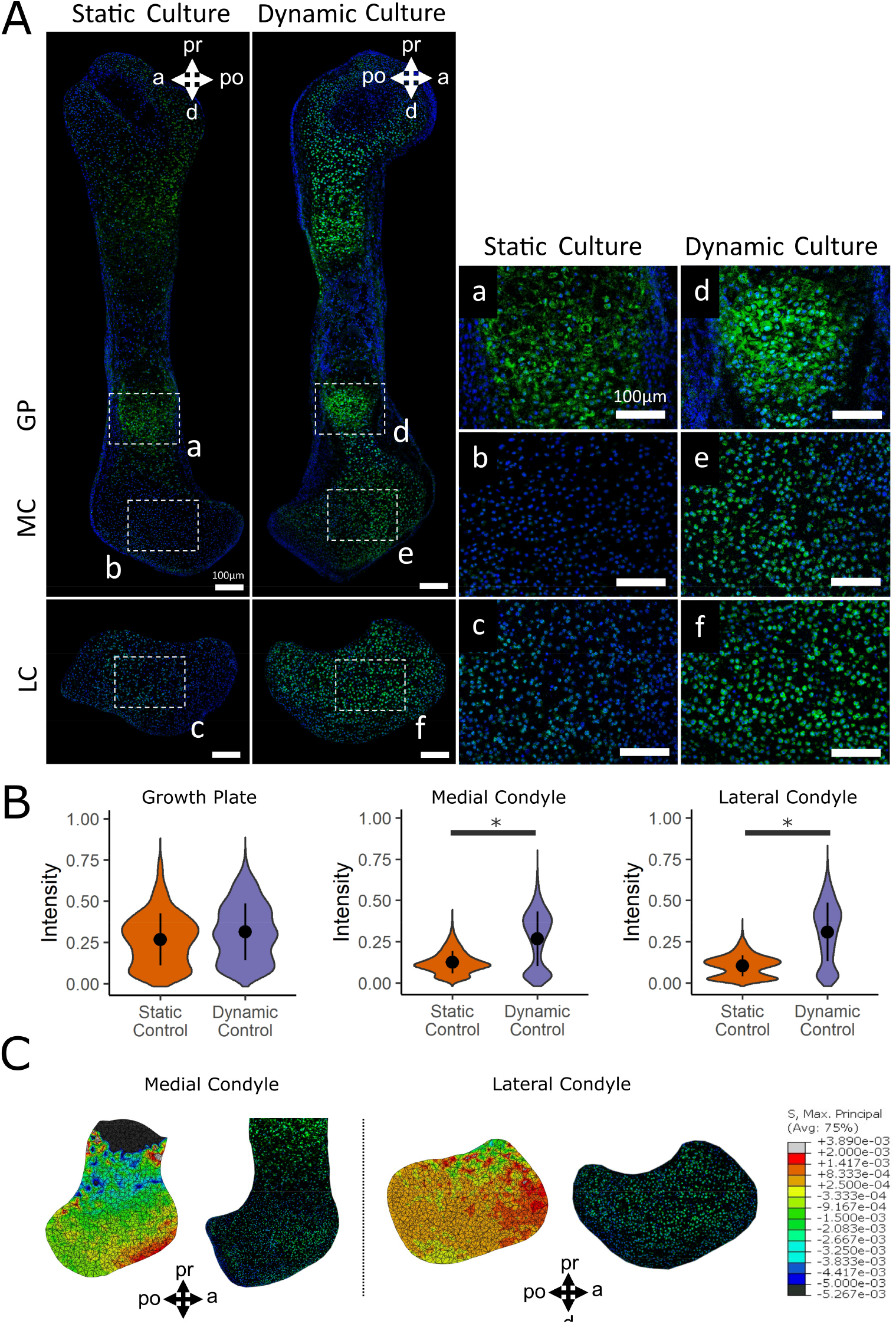
TRPV4 protein expression is mechanically regulated in developing joint cartilage and localized to regions of high maximum principal stress. (A) Immunofluorescence staining of the TRPV4 protein in the growth plate (GP, a, d), medial condyles (MC, b, e) and lateral condyles (LC, c, f) of femora subjected to static and dynamic culture conditions (n=3 limbs per group). Blue; DAPI, Green; TRPV4. Scale bars = 100 µm. (B) Quantification of pooled cell TRPV4 protein immunofluorescence intensity within the growth plate, medial condyle and lateral condyle. * indicates a significant difference between groups (p<0.05). (C) Predicted spatial patterns of maximum principal stress in the medial (left) and lateral (right) condyles from finite element models of dynamically cultured mouse hindlimb distal femora with equivalently oriented sections stained for TRPV4. pr; proximal, d; distal, a; anterior, po; posterior.

To further interrogate the association between biophysical stimuli and TRPV4 protein expression, we examined if TRPV4 preferentially localized to sites within the femoral condyles that experience high stress during loading. A 3D finite element (FE) model of the dynamically loaded limb explants was constructed to compute the stress distributions within the distal femur. Maximum principal stresses were highest in the anterior aspect of the lateral condyle and lowest in the hypertrophic region adjacent to the mineralized cartilage (Fig 3C). Stresses were generally higher in the lateral condyle than in the medial condyle, potentially correlating with increased mechanically-mediated growth of the lateral condyle than the medial (Fig 2). Regions of high maximum principal stresses in the femoral condyles co-localised with regions of increased TRPV4 intensity (Fig 3C). Maximum principal stresses were orders of magnitude higher at the anterior-distal aspect of the medial condyle compared to the rest of the condyle, which corresponded to higher TRPV4 protein expression in the anterior region relative to the rest of the condyle (Fig 3C). In the lateral condyle, a pronounced reduction in maximum principal stress in the distal prominence of the lateral condyle relative to the rest of the condyle was associated with reduced TRPV4 intensity in this region compared to the rest of the condyle (Fig 3C). Co-localisation between high maximum principal stresses and elevated TRPV4 protein expression was not observed in the hypertrophic region, as although maximum principal stresses were very low in the hypertrophic region due to stress shielding from the mineralised cartilage (Fig 3), TRPV4 protein expression was very high in the hypertrophic chondrocytes. The lack of difference in TRPV4 protein expression between dynamically and statically cultured growth plates indicate that TRPV4 in hypertrophic chondrocytes may potentially be regulated by non-mechanical pathways or by cell- or tissue-intrinsic mechanical signals. Altogether, these findings indicate that the spatial variation of biophysical stimuli generated by dynamic loading *ex vivo* drives local concentrations of TRPV4 in the cartilaginous epiphysis, but not at the growth plate.

### Joint cartilage growth and shape are mediated by TRPV4 activity

Having shown that murine joint cartilage growth and TRPV4 protein expression are mechanically regulated, we next set out to determine if loading-induced joint cartilage growth is dependent on TRPV4 activity. A TRPV4 specific antagonist, RN-1734, was diluted in culture media to block TRPV4 activity (Gilchrist et al., 2019) in statically and dynamically cultured limb explants. One hindlimb from each animal was cultured with 10µM RN-1734 while the contralateral limb was cultured with the drug vehicle (DMSO) alone.

Administering the TRPV4 antagonist in static cultures had minimal effects on cartilage growth and shape, with no identifiable differences in contours between the blocked and control static groups (Fig 4B and 4C), and only one measurement (tibial width, p<0.05) showing significant differences (Fig 4B and S1.2). In contrast, the inhibition of TRPV4 in dynamic limbs significantly suppressed the growth of six out of eight cartilage joint features (*p*<0.05, Fig 4A and S1.3), and joint shapes of the limbs resembled those of static distal femora (Fig 4C). Specifically, the posterior curl protrusions in the dynamic blocked limbs were subtle in comparison to dynamic controls (Fig 4C, *), and the outward expansion of the lateral condyle seen in the dynamic controls (Fig 4C, †) was absent in the dynamic blocked group. Medial condyle width and height were not significantly affected by exposure to the antagonist (Fig 4A). The larger effect of loading on growth of the lateral condyle relative to the medial condyle is in line with the higher maximum principal stresses observed in the lateral, compared to the medial, condyle of our FE model. Therefore, blocking TRPV4 in static cultures has minimal effects on joint cartilage growth and shape, while blocking TRPV4 in dynamic limbs almost eliminates the growth-enhancing effects of mechanical loading on the joint cartilage, demonstrating the specific mechanoregulatory role of TRPV4 in limb development. With regards to diaphyseal mineralization, blocking TRPV4 in statically cultured limbs led to a significant (p<0.05) decrease in tibial mineralization (Fig S2), however this effect was not found in femoral mineral. In addition, blocking TRPV4 in dynamically cultured limbs did not affect femoral or tibial mineralization (Fig S2). Thus, it is unlikely that TRPV4 activity significantly influences mineralization in our culture model.

**Figure 4:**
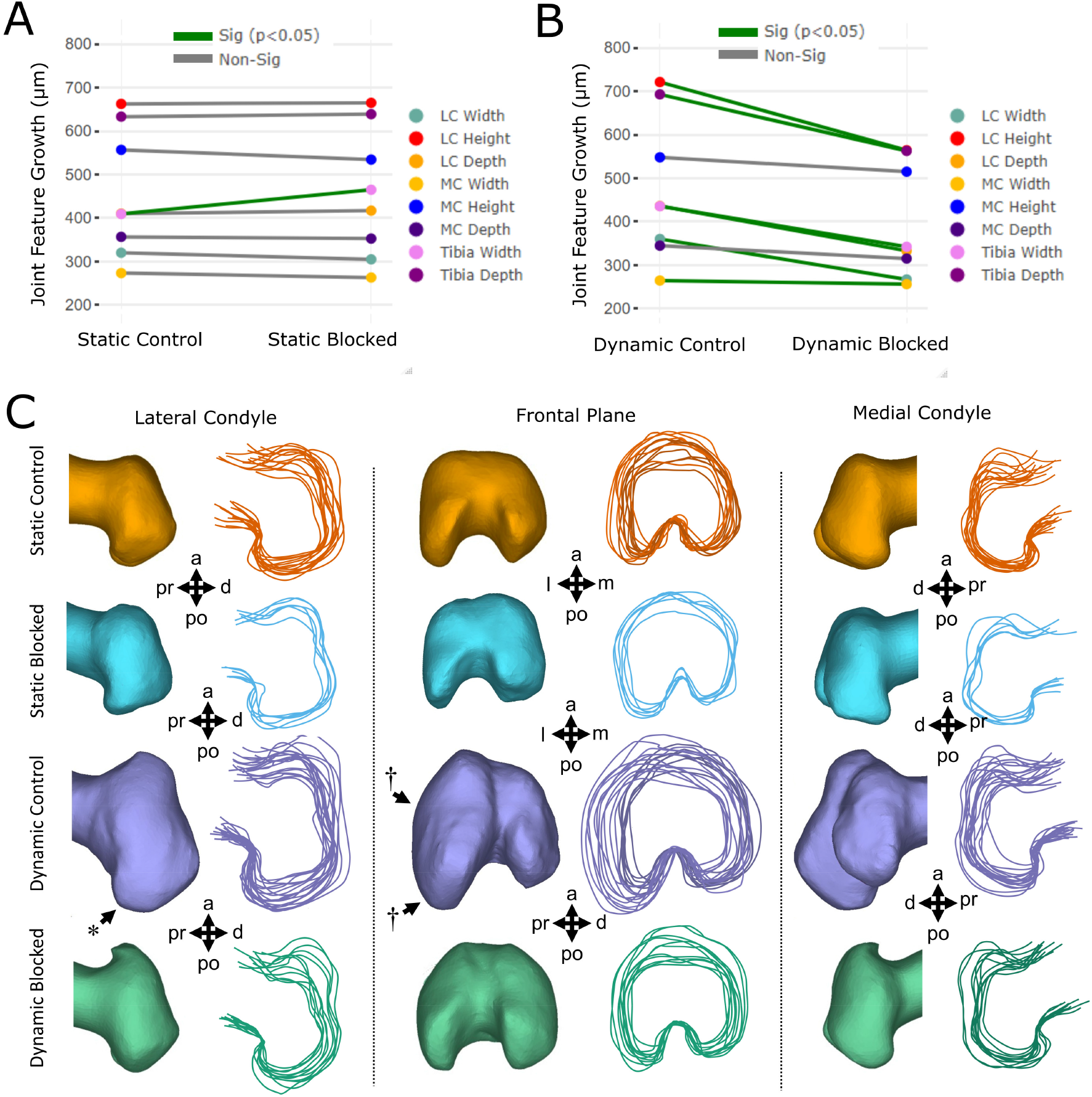
TRPV4 mechanotransduction is crucial for loading-induced joint cartilage growth and shape in dynamically cultured limbs. (A) Average paired differences in growth for joint cartilage features in static vehicle control limbs versus static cultured limbs exposed to TRPV4 antagonist (n=6 limbs per group). Green lines; significant paired feature differences (p<0.05). Grey lines; non-significant feature differences. (B) Average paired differences in growth for joint cartilage features (each line represents one of eight features) in dynamic vehicle control limbs versus dynamic cultured limbs exposed to TRPV4 antagonist RN-1734 (n=12 per group). Green lines; significant paired feature differences (p<0.05). Grey lines; non-significant feature differences. (C) Joint shape contours for distal femora cultured in static and dynamic conditions, exposed to TRPv4 antagonist RN-1734 (10µM) or vehicle control only. n=12 per group. * represents more prominent posterior curl observed in dynamically loaded limbs. † represents larger lateral expansion found in the lateral condyle of dynamically loaded limbs. pr; proximal, d; distal, a; anterior, po; posterior, m; medial, l; lateral.

Statically and dynamically cultured limbs exposed to RN-1734, or the drug vehicle, were stained for the TRPV4 protein to further validate that TRPV4 protein expression is directly mechanoregulated and elucidate whether it is potentially self-regulated. Blocking TRPV4 activity in static cultures had no significant effects on regional TRPV4 protein expression (Fig. 5A) or TRPV4 intensity (Fig. 5B) in the condyles, or in the growth plate (Fig. S2a). However, while TRPV4 protein expression in dynamic blocked condyles was still significantly higher than that seen in the static control and static blocked limbs (Fig 5B), expression in the condyles was significantly reduced when compared to dynamic controls (Fig. 5A and 5B), indicating that TRPV4 mechanotransduction upregulates TRPV4 expression in joint cartilage. In contrast, blocking TRPV4 activity in dynamic cultures had no significant effects on intensity in the growth plates (Fig. S2b). We conclude that TRPV4 protein expression and function is associated with loading-induced growth and shape in prenatal joint cartilage, and that TRPV4 activity in the growth plates may be differentially regulated.

**Figure 5:**
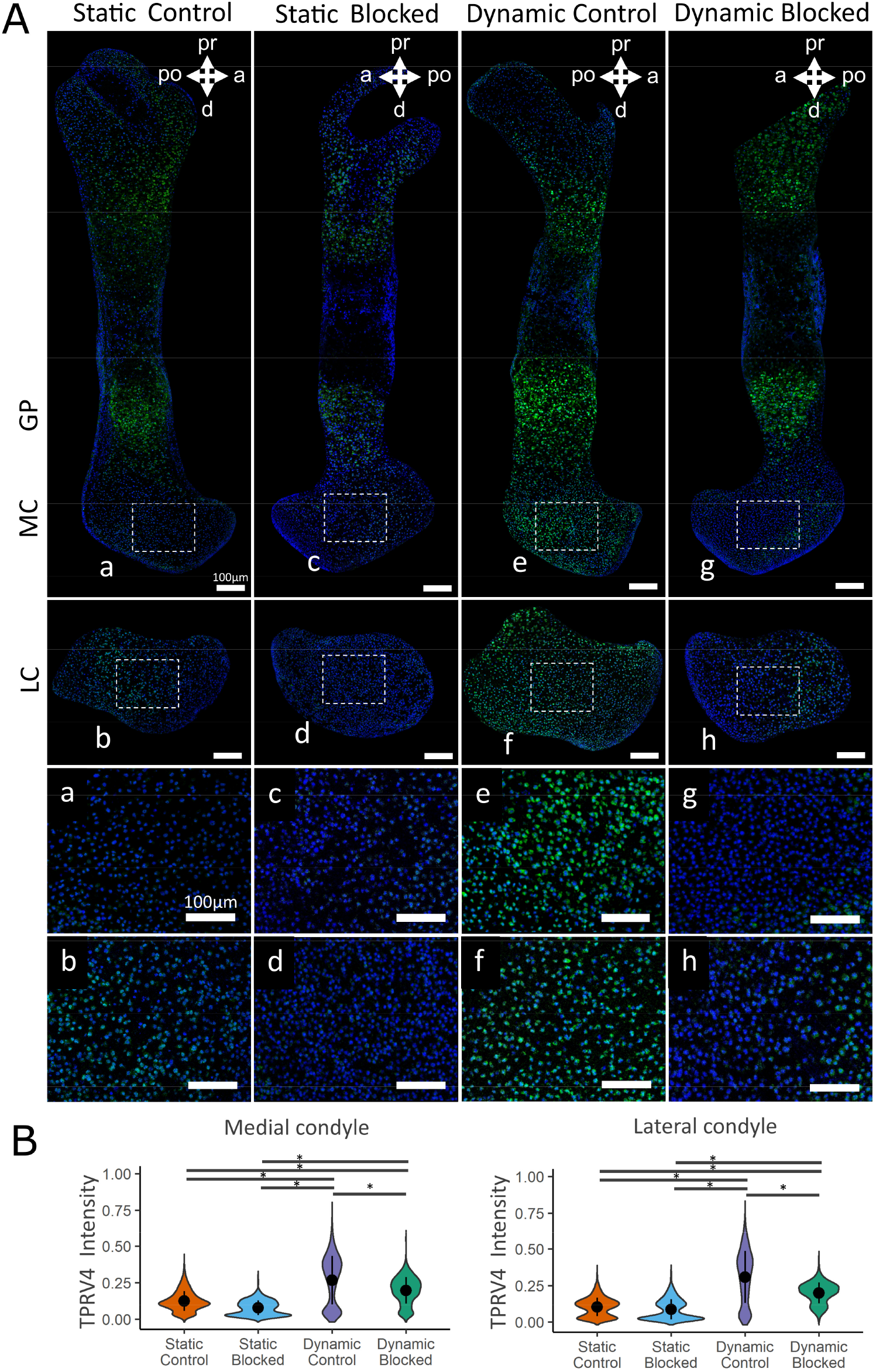
TRPV4 activity is essential for the mechanoregulation of TRPV4 expression in dynamically cultured limbs. (A) Immunohistochemical staining of TRPV4 protein in the medial condyle (a, c, e and g) and lateral condyle (b, d, f and h) of contralateral limbs subjected to static or dynamic culture conditions with the TRPV4 antagonist RN-1734 or vehicle control only (n=3 limbs per group). Blue; DAPI, Green; TRPV4. (B) Quantification of pooled cell TRPV4 protein immunofluorescence intensity within the femoral condyles of static and dynamic limbs exposed to TRPV4 antagonist or vehicle control only. Black bars with * indicate a significant difference between groups (p<0.05). GP; growth plate, MC; medial condyle, LC; lateral condyle, pr; proximal, di; distal, an; anterior, po; posterior.

### TRPV4 mediates chondrocyte proliferation and matrix synthesis in response to loading during skeletal development

Next, we sought to determine the specific cell activities regulated by TRPV4 that drive joint development and morphogenesis. We investigated the involvement of TRPV4 activity with cell proliferation and matrix synthesis, key contributors to skeletal growth and morphogenesis (Brunt et al., 2016; Decker et al., 2017). Cultured limbs were sectioned and labelled for cell mitosis with phosphohistone H3 (PHH3) using immunofluorescence techniques. The proportion of proliferating cells was calculated by normalising to the total number of cells in the medial and lateral condyle regions. Other sections were histologically stained for proteoglycans using Alcian blue and for collagen using picrosirius red.

Subjecting the limbs to dynamic culture resulted in a 4-fold significant increase in the proportion of proliferating chondrocytes in both the lateral condyle and medial condyle (*p*<0.05, Fig. 6) compared to static culture. Notably, the proportion of proliferating cells was 2-fold higher in the lateral condyle compared with the medial condyle in loading conditions, in line with the greater changes in mechanically-mediated growth observed in the lateral condyle compared to the medial condyle. Under static culture conditions, blocking TRPV4 led to a modest decrease in proliferation, which was only significant in the lateral condyle (*p*<0.05, Fig. 6). Introducing the TRPV4 antagonist to dynamic limbs suppressed the mechanically-induced response in both lateral and medial condyles (*p*<0.05, Fig. 6) to levels equivalent to static controls. Therefore, loading-induced proliferation in joint cartilage is strongly mediated by TRPV4 activity.

**Figure 6:**
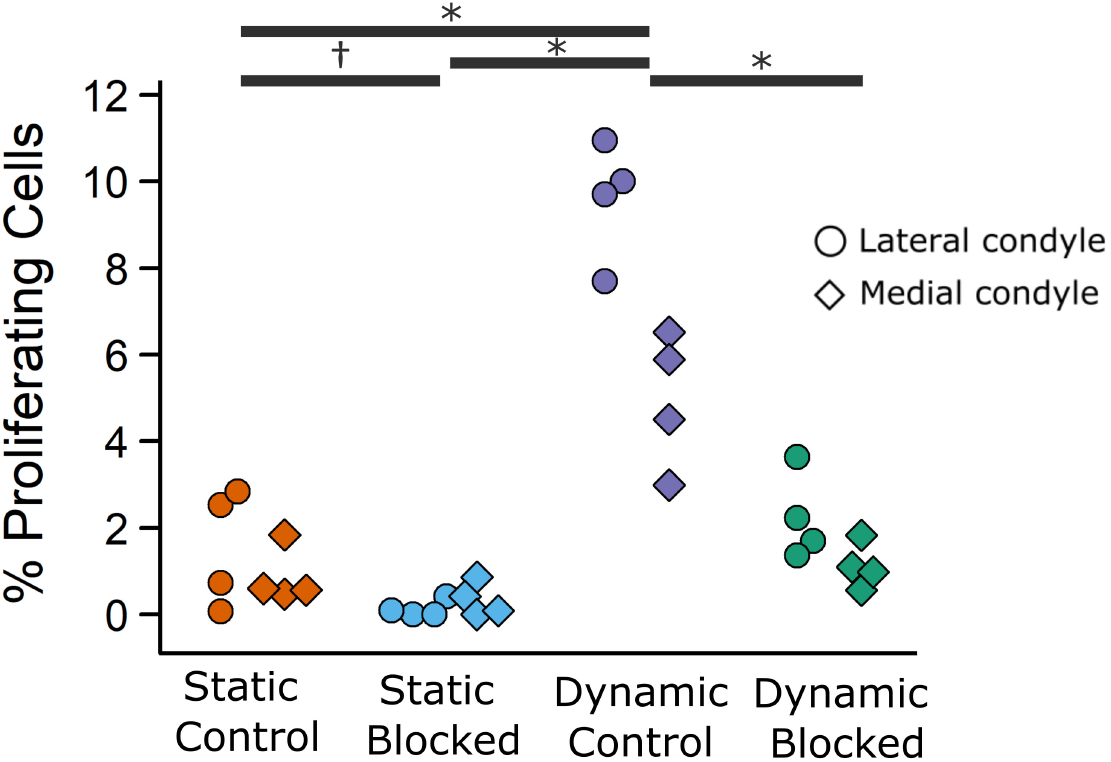
TRPV4 mechanoregulates loading-induced cell proliferation in the femoral condyles of dynamically cultured limbs. Figure illustrates the proportion of proliferating cells within the lateral and medial condyle regions, where each point represents an ensemble average of three sections from each sample (n=4 limbs per group). * indicates significant difference (p<0.05) between groups for the lateral and medial condyle. † indicates significant difference (p<0.05) between groups for the lateral condyle only.

In terms of matrix deposition, statically cultured control limbs displayed light proteoglycan (Fig. 7) and collagen (Fig. 8) staining in the condyles, which appeared unaffected when TRPV4 activity was blocked. Dynamic stimulation in control cultures potently upregulated matrix synthesis, as both proteoglycans (Fig. 7) and collagen (Fig. 8) matrix appeared consistently denser throughout the femoral condyles as compared to static cultures. Blocking TRPV4 reduced proteoglycans in dynamic cultures to levels observed in static control samples (Fig. 7), while collagen deposition appeared only partially reduced in blocked dynamic cultures compared to control dynamic cultures (Fig. 8). Together, these data suggest that TRPV4 mechanotransduction in prenatal joints involves the activation of the chondrocyte proliferation and matrix biosynthesis pathways as part of the anabolic response to dynamic loading.

**Figure 7:**
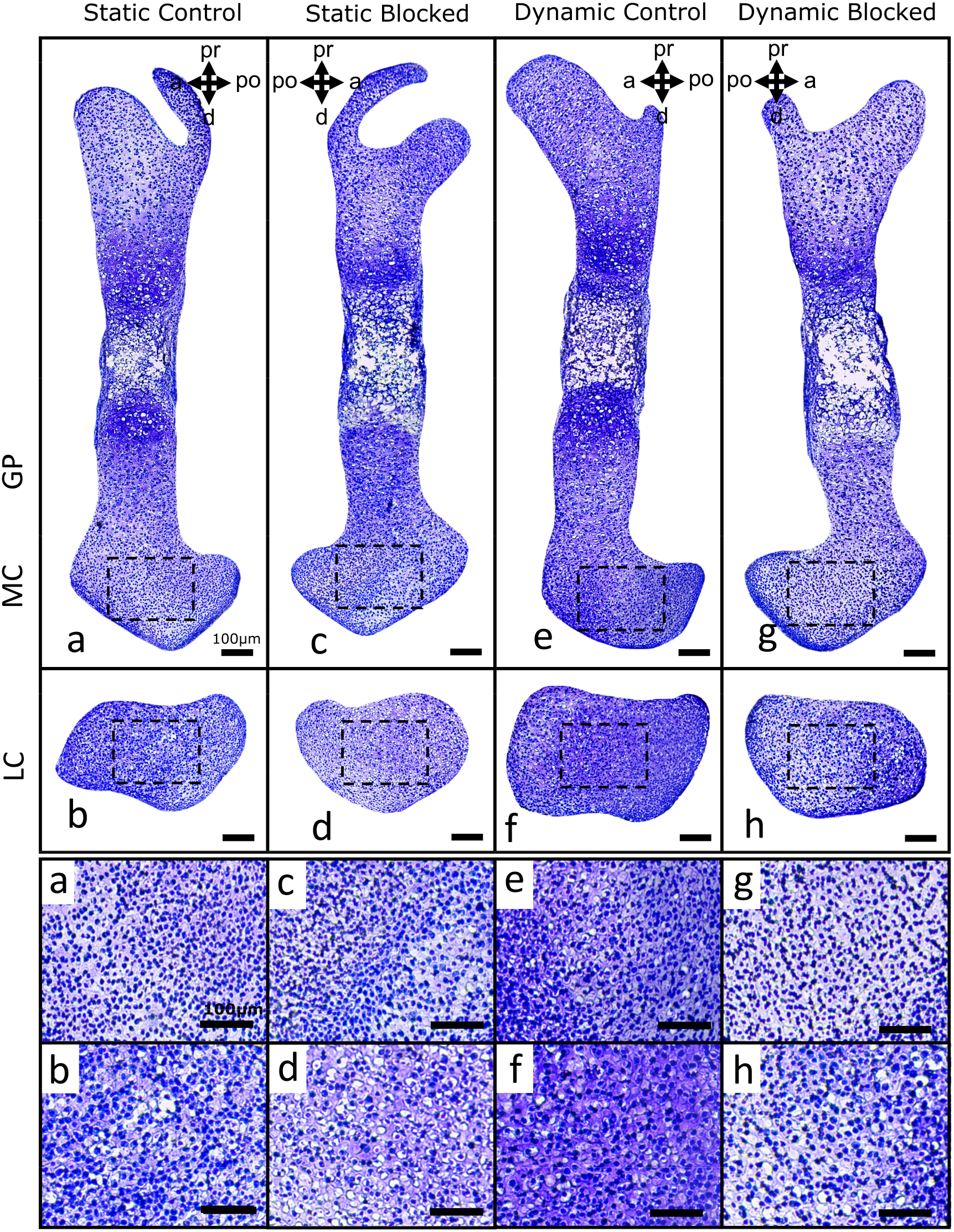
TRPV4 activity mediates loading-induced proteoglycan synthesis in the femoral condyles of dynamically cultured limbs. Proteoglycan deposition (histologically assessed using toluidine blue stain) in the medial condyles (a, c, e, g) and lateral condyles (b, d, f, h) of femora subjected to static or dynamic culture, with TRPV4 antagonist RN-1734 (10µM) or drug vehicle (DMSO) (n=3 limbs per group). Boxes represent areas expanded for figures a-h. GP; growth plate, MC; medial condyle, LC; lateral condyle, pr; proximal, di; distal, an; anterior, po; posterior.

**Figure 8:**
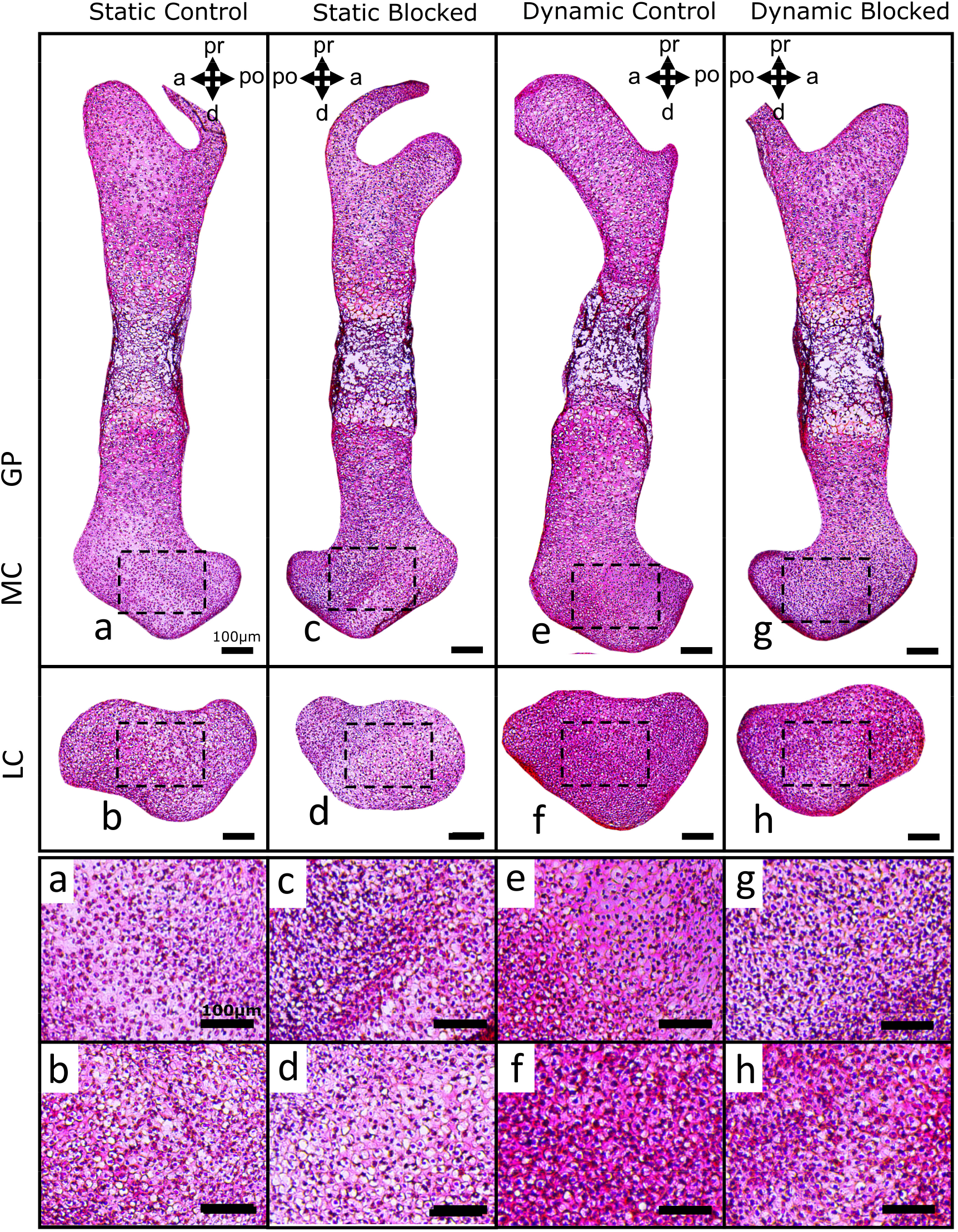
TRPV4 mechanotransduction partially regulates loading-induced collagen deposition in the femoral condyles of dynamically cultured limbs. Collagen deposition (histologically assessed using picrosirius red) in the medial condyles (a, c, e and g) and lateral condyles (b, d, f and h) of femora subjected to static or dynamic culture, with TRPV4 antagonist RN-1734 (10µM) or drug vehicle (DMSO) (n=3 limbs per group). Boxes represent areas expanded for figures a-h. GP; growth plate, MC; medial condyle, LC; lateral condyle, pr; proximal, di; distal, an; anterior, po; posterior.

## DISCUSSION

Previous studies have identified the importance of TRPV4 in cartilage and bone mechanotransduction, and separately, the involvement of TRPV4 channelopathy in skeletal malformations, yet its role in the regulation of skeletal development has not previously been investigated. In this study, we first validated a novel model where we demonstrate that mechanically loading mouse embryonic limb explants *ex vivo* stimulates joint cartilage growth and morphogenesis, but not diaphyseal mineralization. We next revealed that TRPV4 protein is present in developing cartilaginous tissues and is spatially localized to sites of high biophysical stimuli, as indicated by our computational model of joint loading. We then demonstrated that TRPV4 protein expression is mechanically self-regulated and that regulatory pathways initiated by TRPV4 are essential for the anabolic cartilage response to physical stimuli. Finally, we show that this mechanoregulatory role of TRPV4 in skeletal development is achieved via the control of cell proliferation and matrix biosynthesis. Since TRPV4 was co-localized to sites of high biophysical stimuli, we speculate that local stress patterns lead to upregulation of TRPV4 in the joint tissues, leading in turn to increased proliferation, matrix deposition and consequent regional growth. This indicates a mechanism by which mechanical loading could direct morphogenesis during cartilage development, and by which TRPV4 channelopathies may lead to joint malformations (Nilius & Voets, 2013). Therefore, while the involvement of TRPV4 in skeletal development is evidenced by the impact of TRPV4 channelopathies on abnormal skeletal formation (Nilius & Voets, 2013), this study demonstrates the specific mechanoregulatory involvement of TRPV4 in prenatal joint development.

A compelling aspect of our findings is that they reflect what have been found in cell culture studies at the cell level, but uniquely, at cell, tissue and organ levels in whole limb explants. Previous studies using chondrocyte-laden constructs, cultured MSCs or chondroprogenitors found a similar dependence on TRPV4 in terms of loading induced upregulation of cartilage matrix gene expression markers *ACAN, COL2α1*, as well as proteoglycans and collagen matrix accumulation (Corrigan et al., 2018; O’Conor et al., 2014; Willard et al., 2021). Furthermore, exposing MSCs and chondrocytes to TRPV4-specific agonists without mechanical stimulation leads to enhanced matrix synthesis at similar levels to that found in mechanically loaded cells (Corrigan et al., 2018; Willard et al., 2021). Together, the results from these studies combined with ours provide evidence for a critical regulatory role for TRPV4 in cartilage growth, and highlights potential therapeutic targets for modulating cartilage properties. Furthermore, since our findings indicate that TRPV4 localizes to regions of high mechanical stimuli, it may be possible to use mechanical stimulation combined with agonists to direct the effect of TRPV4-mediated growth to specific regions and modulate cartilage shape. Mechanically mediated collagen deposition was only partially suppressed when TRPV4 was blocked, possibly indicating that synthesis of only some collagen types are regulated by TRPV4, since picrosirius red stains for all collagens. Immunofluorescence imaging of specific collagen networks such as collagen II that emerge during cartilage development (Ahmed & Nowlan, 2020) may provide further insight into the influence of TRPV4 on collagen synthesis and organisation.

The positive effect of dynamic loading on chondrocyte proliferation in the femoral condyles was abolished by blocking TRPV4. While not previously investigated in chondrocytes, these findings are consistent with previous work that observed increased proliferation of micromass iPSC-derived chondroprogenitors treated with the TRPV4 agonist GSK101 (Willard et al., 2021). Blocking ion channel dependent Ca^2+^ signalling inhibits mechanically induced parathyroid hormone-related protein (PTHrP) upregulation in chondrocytes exposed to cyclic mechanical strain (Tanaka et al., 2005). Therefore, the PTHrP / Indian hedgehog (IHH) signalling loop which regulates chondrocytic proliferation and matrix synthesis (Amizuka et al., 1994; Tanaka et al., 2005) could be mediated by TRPV4. Our results combined with those of others suggests a functional role further upstream of genes such as *SOX9* and *TFG-β3*, key markers and regulators of chondrogenesis (Bi et al., 1999; DeLise et al., 2000). Controlled modulation of TRPV4 may therefore be an attractive direction for enhancing the properties of developmentally inspired tissue engineered constructs that aim to recapitulate the processes of cartilage and bone development (Freeman & McNamara, 2017; Lenas et al., 2009). Interestingly, while TRPV4 protein expression was regulated by mechanical loading in the joint cartilage, expression in the hypertrophic cartilage appeared to be consistently high and unaffected by mechanical loading. Hypertrophic chondrocytes swell to a larger cell volume when transitioning from resting chondrocytes (Kronenberg, 2003), proposed to be a critical determinant of rudiment lengthening (Bush et al., 2008), therefore we speculate that increased cell membrane tension due to hypertrophy may be intrinsically upregulating TRPV4 protein expression.

A limitation encountered in this study was that our culture model was not suitable for investigation into the involvement of TRPV4 in diaphyseal mineralization. Endochondral ossification is influenced by mechanical stimulation *in utero*, consistently in the chick (Nowlan et al., 2008) but only in some rudiments in the mouse (Nowlan et al., 2010). Dynamic culture did not affect mineralization in the murine limb explants as it has previously in our chick hindlimb explant model (Khatib et al., 2021), likely due to the reliance of endochondral ossification on blood vessel invasion in the mammalian limb (Kronenberg, 2003; Nowlan et al., 2007). Previous groups have developed tissue constructs mimicking aspects of endochondral ossification (Freeman & McNamara, 2017; Lenas et al., 2009), which may be a more viable route to investigating the role of TRPV4 in bone development *in vitro*. Previous work has demonstrated a potent efficacy of RN-1734 of inhibiting TRPV4-induced Ca^2+^ signaling in cultured MSCs, showing a similar effect to other antagonists including the widely used GSK205 (Gilchrist et al., 2019). Our findings are in line with previous work from our group looking at the effects of gadolinium on chick cartilage morphogenesis (Parisi et al., 2018), which further substantiates our results in the mouse model. Considerable off-target effects due to RN-1734 seem unlikely, due to the absence of effect on growth, matrix deposition and proliferation in static limbs.

In conclusion, the results from this study indicate that TRPV4-mediated mechanotransduction is crucial for the mechanical regulation of cartilage development during skeletogenesis, partially through the upregulation of cell proliferation and matrix synthesis pathways. As a potent regulator of cartilage formation, TRPV4 may be a valuable target for the development of therapeutic disease modifying drugs, or developmentally-inspired tissue engineering strategies for skeletal tissue repair.

## METHODS

### Limb culture

Mouse (strain C57/B16) embryos were harvested at E15.5 (Theiler stage 24). Hindlimbs were dissected from the spine and the bulk of soft tissue removed under the microscope to improve culture media and drug penetration. Limbs were cultured in vitro at an air-liquid interface with dynamic loading within a mechanostimulation bioreactor or in static conditions within a petri dish over a 6-day period as previously described (Chandaria et al., 2016).

All culture vessels were filled with osteogenic media consisting of basal media (αMEM with GlutaMAX supplement), supplemented with 1% pen/strep and amophotericin B, 100µM ascorbic acid, 2mM β-glycerophosphate and 100nM Dexamethasone. TRPv4 mediated Ca^2+^ activity (Gilchrist et al., 2019) was blocked using RN-1734 (Sigma Aldrich) dissolved in DMSO (Sigma Aldrich), diluted into the complete basal media at a final concentration of 10µM as used previously (Gilchrist et al., 2019). The same volume of DMSO was diluted in basal media for the vehicle control.

### Mechanical stimulation

For the dynamic culture condition, the bioreactor was programmed to apply 2mm sinusoidal compressive displacements to the foam supports with limbs at 0.67 hertz, which induced a sagittal knee flexion of approximately 14° (+/- 2°) (Fig. 1A). Within each 24-hour period, three 2-hour intervals of mechanical stimulation with 6-hour periods of rest in between. For the static culture conditions, the foam supports with limbs were placed in an enclosed petri dish.

Cultured left and right hindlimbs served as contralateral pairs for three paired comparison culture groups: (1) static culture vs dynamic culture, (2) dynamic culture with RN-1734 vs dynamic with DMSO (vehicle control), or (3) static with RN-1734 vs static with DMSO (Fig 1B).

### Cartilage and mineral measurements and shape analysis

To quantify cartilage growth and mineral length, measurements were taken from 3D representations of the hindlimb knee joints generated through optical projection tomography (OPT) techniques optimised for embryo imaging (Quintana et al., 2009). Prior to OPT, samples were dehydrated in ethanol, stained with 0.055% Alcian blue for 5 hours, cleared in 1% KOH for 2 hours, and then stained with 0.01% alizarin red for 2 hours at room temperature, permitting selective visualisation of the cartilaginous tissues and mineral, respectively. Following OPT scans, image projections were reconstructed (NRecon, Micro Photonics Inc., USA) and segmented into 3-dimensional (3D) models (Mimics 19, Materialise, Belgium) of the cartilage. Eight knee joint cartilage features and two mineral lengths of the 3D models were measured (Fig. 2). For joint feature measurements, the distance between the apexes of anatomical landmarks was collected for each model (3-Matic, Materialise, Belgium). To assess joint feature shape, the cartilage joint models were rigidly registered using N-point registration (3-Matic, Materialise) and the contour of the joints extracted in the medial, frontal and lateral view of the joints. Wilcoxon signed-rank tests with Bonferroni corrections were used to test for paired differences in quantitative cartilage growth variables and mineral length between contralateral limb samples.

### Histology and Immunofluorescence

Cultured hindlimbs were processed in a sucrose gradient (15%, 30%) and embedded in a 30% sucrose and optimal cutting temperature compound (OCT) 50:50 mix. 10µm sections of the medial condyles with full-length femora and lateral condyles were collected along the sagittal axis of the femora and fixed in 4% paraformaldehyde.

For histology, frozen sections were stained with toluidine blue (sulphated proteoglycans stain) for 4 mins and washed with deionised water, or stained in picrosirius red (collagen stain) for 30 mins, followed by 10 mins in acidified water. Sections were imaged using light microscopy with a consistent exposure time (Yenway EX30l; Life Sciences Microscope, Glasgow, UK).

For immunofluorescence, frozen sections were washed with PBTD (0.1% Tween-20 and 1% DMSO in PBS) for permeabilization, blocked with 5% normal goat serum (NGS) for 2 hours, incubated with primary antibodies against TRPV4 protein (1:500, ab39260, Abcam) or Phosphohistone-H3 (1:500, ab5176, Abcam) at 4°C overnight, followed by Alexa Fluor 488-conjugated secondary antibody (1:200, ab150077, Abcam) for 2 hours and DAPI for 3 minutes.

To quantify average regional TRPV4 intensity, immunofluorescence images of the growth plate, and central regions of the medial condyle and lateral condyle were captured at 10x magnification, using sections stained for TRPV4. Edges of the rudiments were not used in the quantification due to high errors in cell segmentation where the density of cell nuclei is highest. Cell nuclei were segmented as primary objects using the DAPI channel to detect chondrocyte locations, and whole cell areas segmented as secondary objects using the TRPV4 channel (488), based on the location of the primary objects. Finally, the average pixel grayscale intensity of Alexa488 (TRPV4) was quantified within the secondary object areas. Significant differences in TRPV4 intensity (pooled across all cells) between culture groups was assessed using the Mann-Whitney U tests for two group comparisons, or one-way ANOVA with Tukey post-hoc tests for four group comparisons.

To quantify the percentage of proliferating cells, images taken at 10x magnification were cropped to the medial and lateral condyle regions only, then cell nuclei areas were segmented using the DAPI stain (CellProfiler™, Broad Institute) and quantified. Number of cells positively expressing PHH3 were manually counted (ImageJ). One way ANOVA with Tukey post-hoc tests were used to assess significant differences between number of proliferating cells between groups.

### Finite element analysis

Principal Stress was calculated through Finite Element Analysis of an ideal limb in ABAQUS. Multiple OPT scans of limbs were collected and separated into distinct components, namely, the distal femur and the bone collar. These components were registered using n-point registration which allowed for average components to be produced. An ideal limb was created using the averaged distal femur and bone collar, with the proximal femur and tibia supplied by the clearest, most representative whole limb OPT scan (S7A). The relative orientation of the femur and tibia were adjusted to match the angle of limbs in the zero position from an image taken at day 6 of culture. A solid spherical joint capsule was then placed surrounding the joint space (S7B). Two boundary conditions were applied to the system. The proximal femur was driven with a ramped displacement designed to match the frequency and amplitude applied to the limbs during *in vitro* mechanical stimulation. The distal tibia was fixed but allowed to freely rotate, reflecting the interaction of the foot with the foam step. All components were provided with linear isotropic elastic properties, with a young’s modulus (MPa) of 1.1 (Tanck et al., 2004), 117 (Tanck et al., 2000) and 0.55 (Nowlan et al. 2012), and a poisons ratio of 0.49 (Tanck et al., 2004), 0.3 (Tanck et al., 2000) and 0.25 (Nowlan et al. 2012), for the cartilage, mineralised region and capsule respectively.

## ACKNOWLEDGMENTS

This work was funded by the European Research Council under the European Union’s Seventh Framework Programme (ERC Grant agreement number 336306)

## Notes

### Competing Interest Statement

The authors have declared no competing interest.

