## Supplementary Figures S1-S7 for "Mechanoregulatory role of TRPV4 in prenatal skeletal development"

### SUPPLEMENTARY MATERIALS

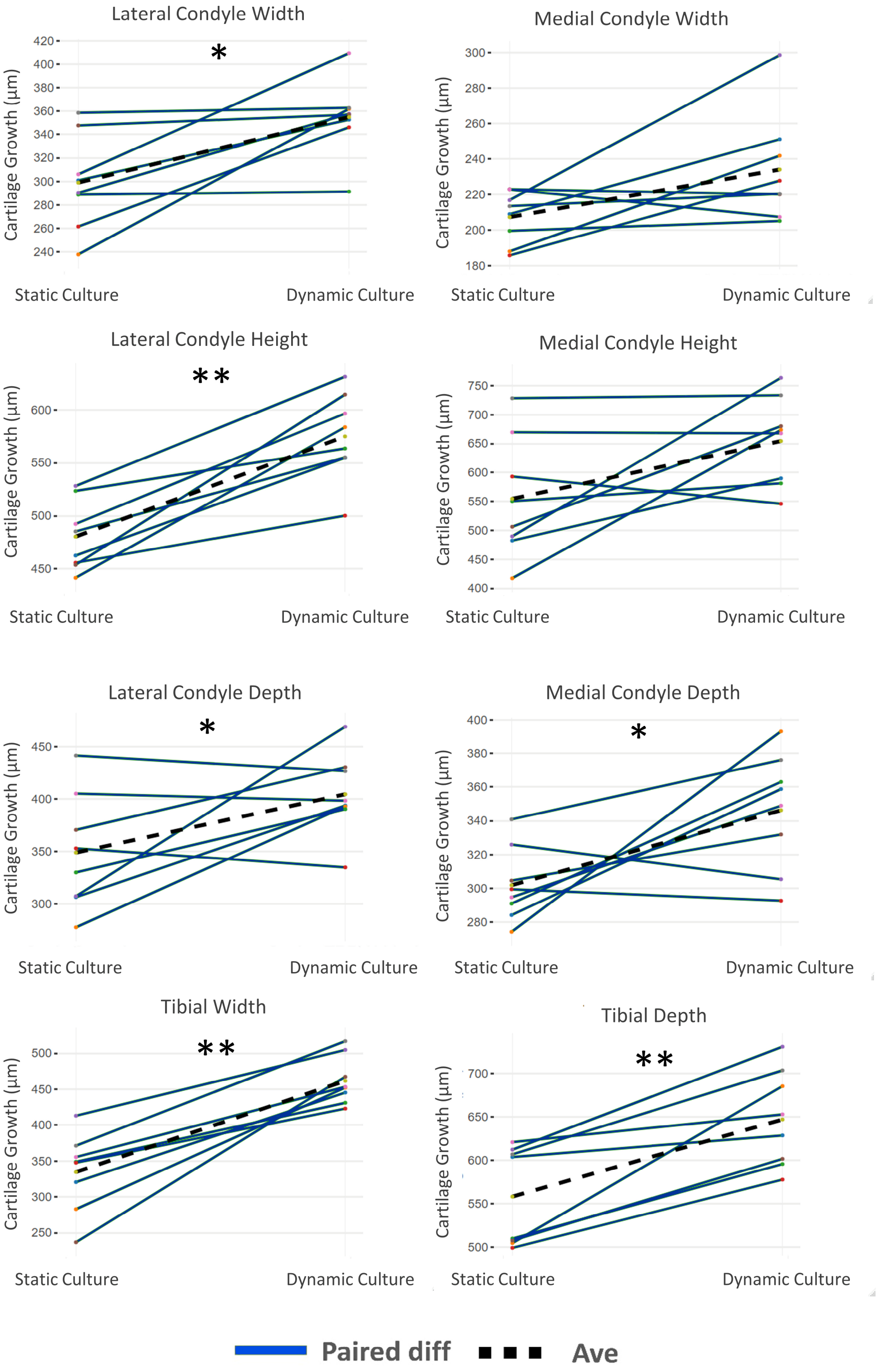


**S1.1 Paired sample differences in joint cartilage feature growth between statically and dynamically cultured limbs.** Each line shows the difference between contralateral limbs of one embryo. * $p<0.05$; ** $p<0.01$; n=8 limbs per group.

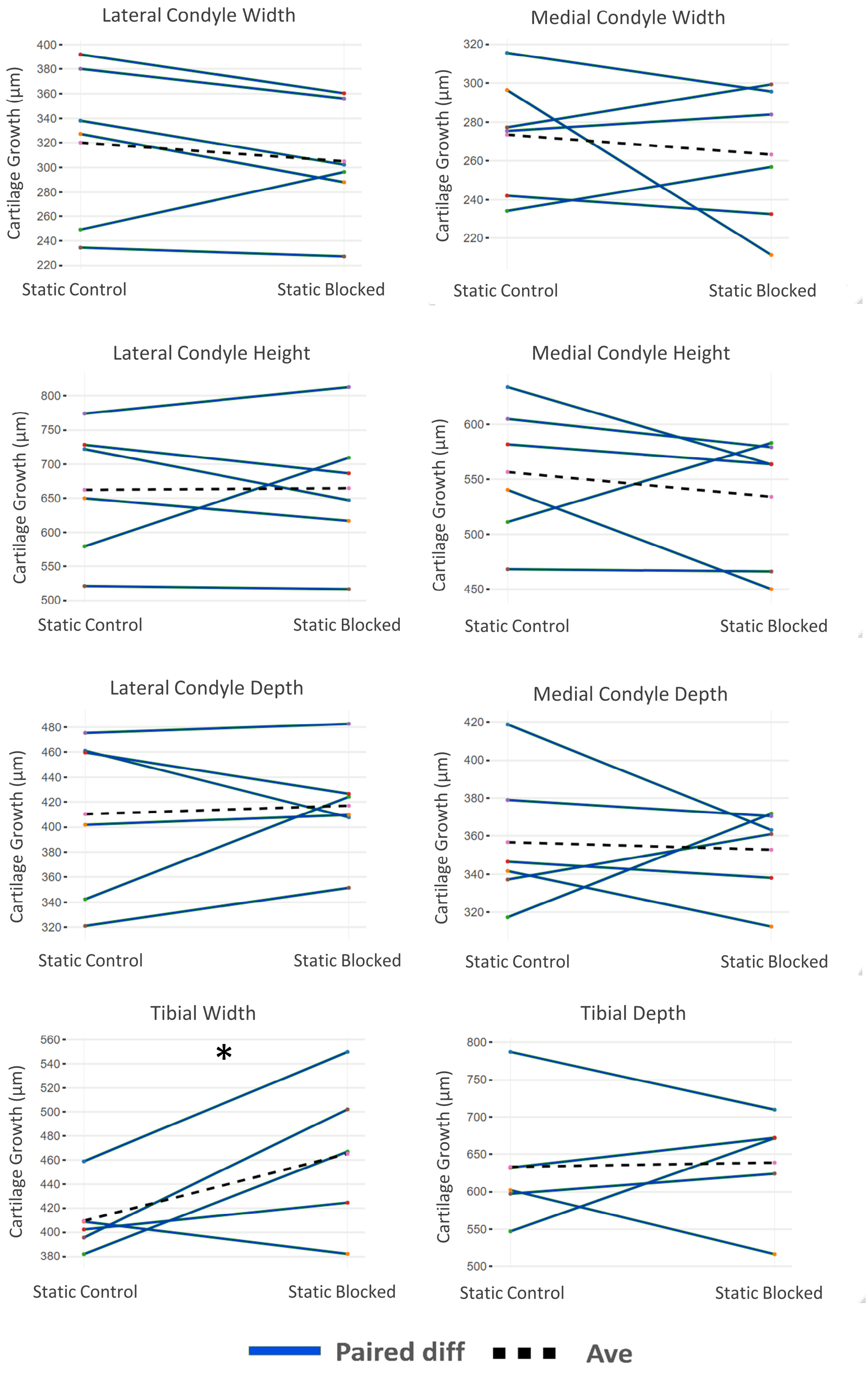


**S1.2 Paired sample differences in joint cartilage feature growth between static vehicle control and static blocked limbs.** Each line shows the difference between contralateral limbs of one embryo. * $p<0.05$; ** $p<0.01$; n=6 limbs per group.

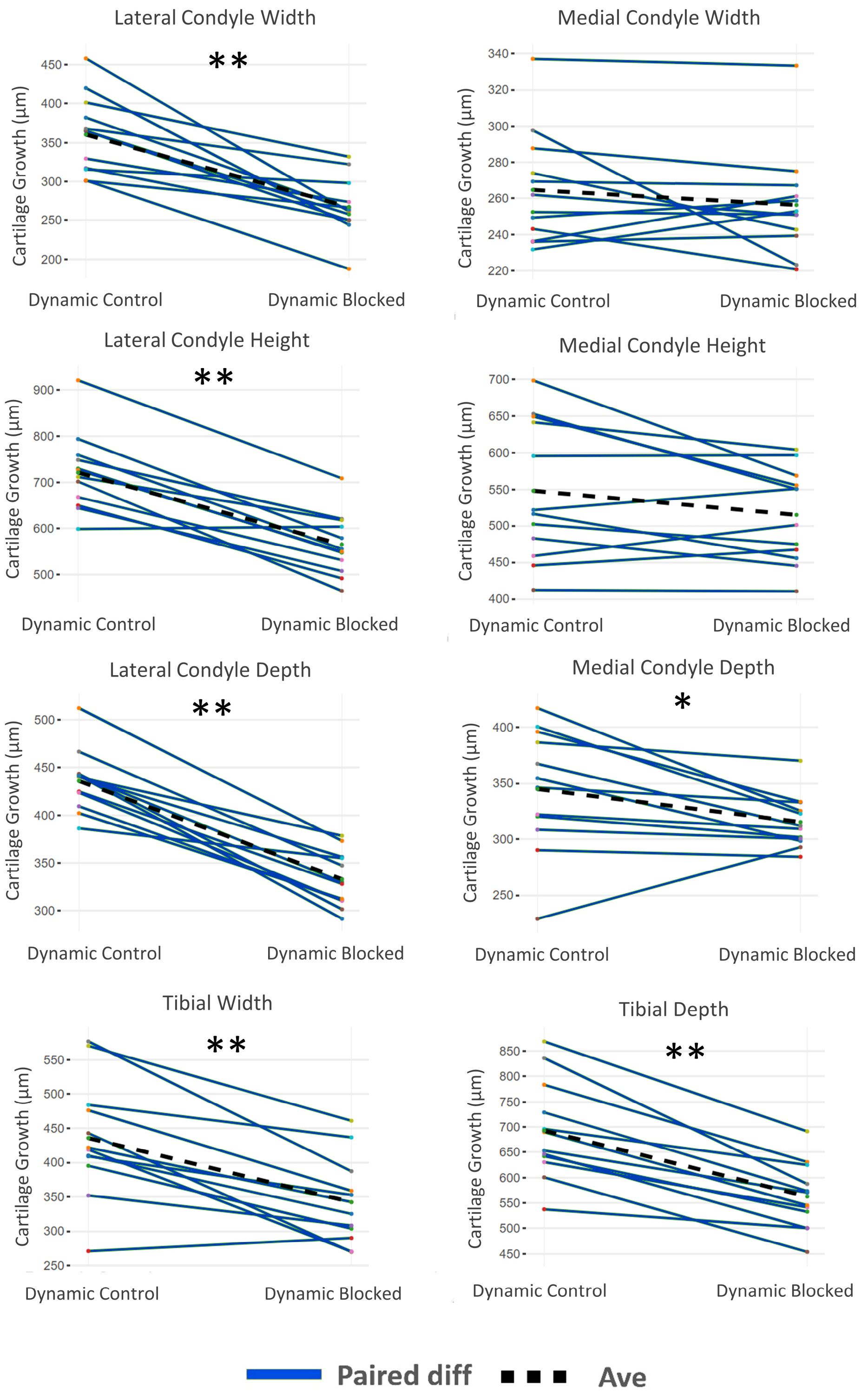


**S1.3 Paired sample differences in joint cartilage feature growth between dynamic control and dynamic blocked limbs.** Each line shows the difference between contralateral limbs of one embryo. * $p<0.05$; ** $p<0.01$; n=12 limbs per group.

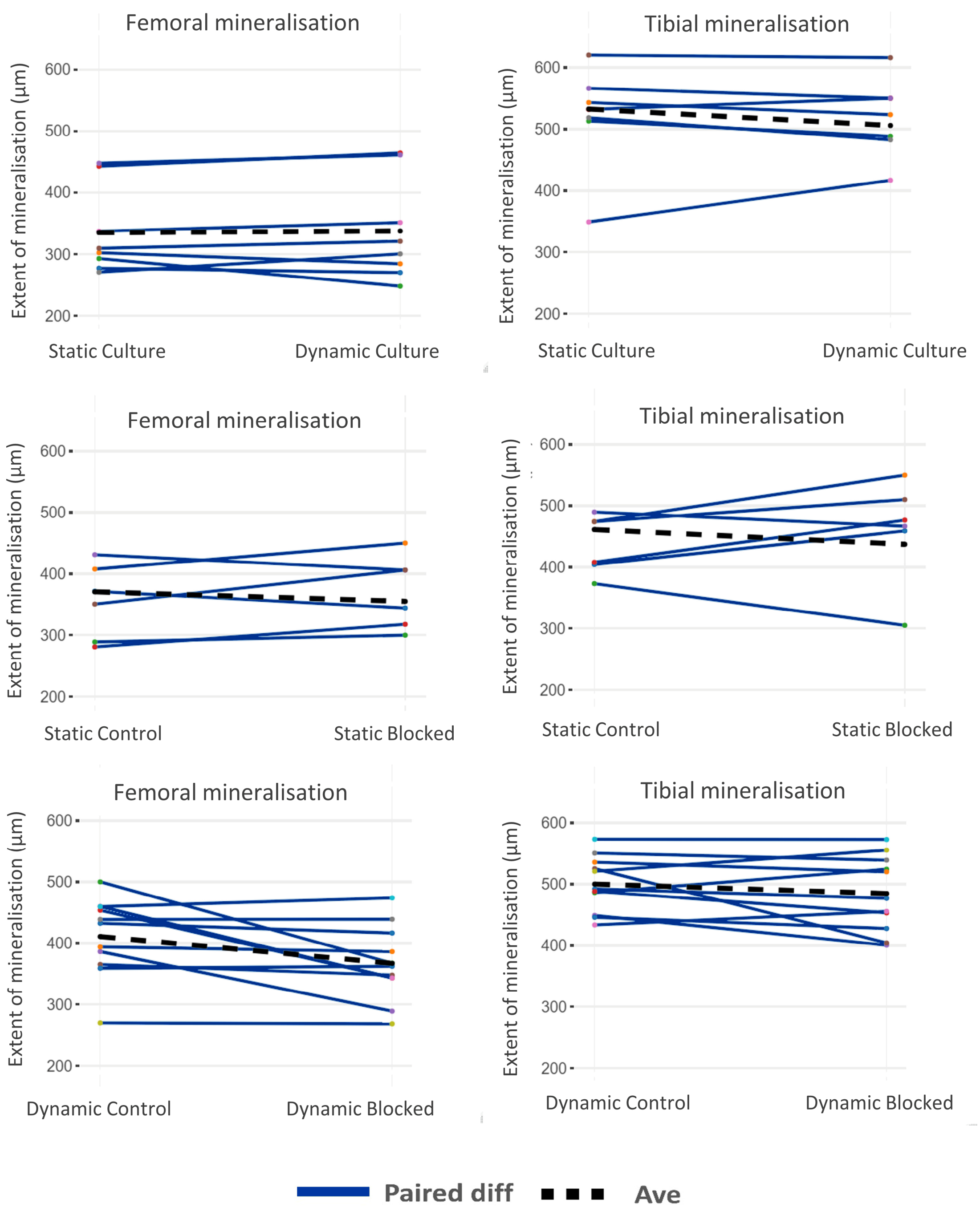


**S2. Paired sample differences in diaphyseal mineralization in all comparison groups.** Each line shows the difference between contralateral limbs of one embryo. * $p<0.05$; ** $p<0.01$; static vs dynamic, n=6; dynamic control vs dynamic blocked, n=10, dynamic control vs dynamic blocked, n=6.

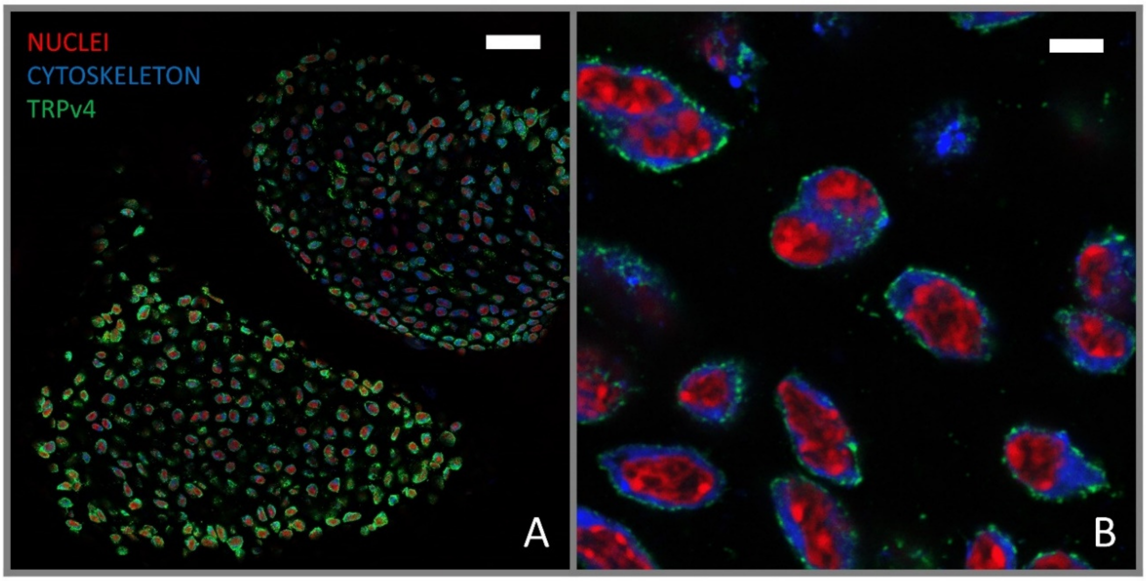


**S3: Immunolocalisation of TRPv4 on the cell membrane of embryonic murine epiphyseal chondrocytes.** Red; nuclei, blue; cell cytoskeleton, green; TRPv4 immunolocalization. Scale bars: 50µm for A and 10µm for B.

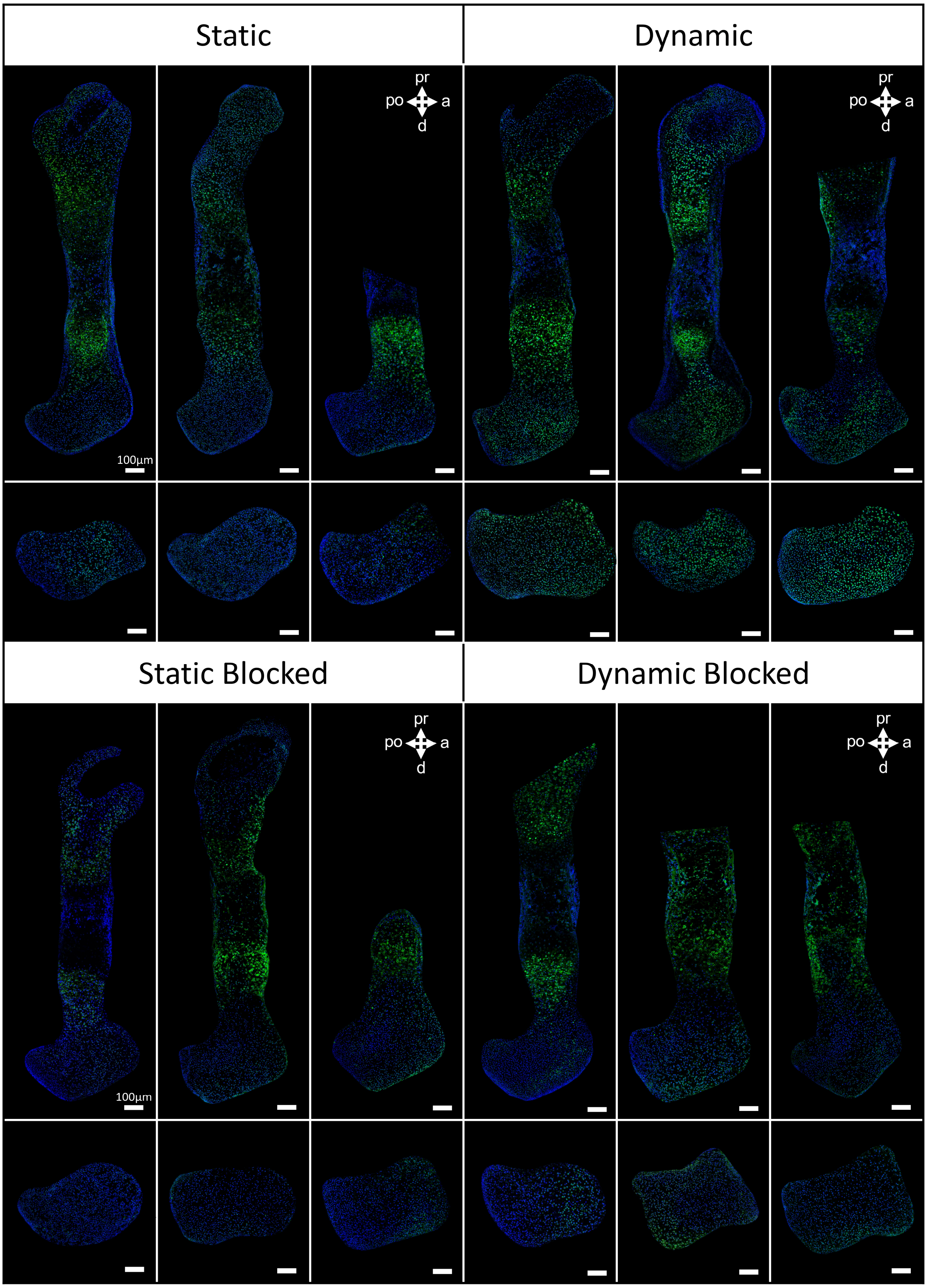


**S4: Regional TRPV4 protein expression within all mouse embryo femora.** Each column represents contralateral limbs from a single embryo split into two comparison groups. The static and dynamic vehicle control limbs (top row) are horizontally mirrored to aid comparison across the whole dataset. n=3 limbs per comparison group.

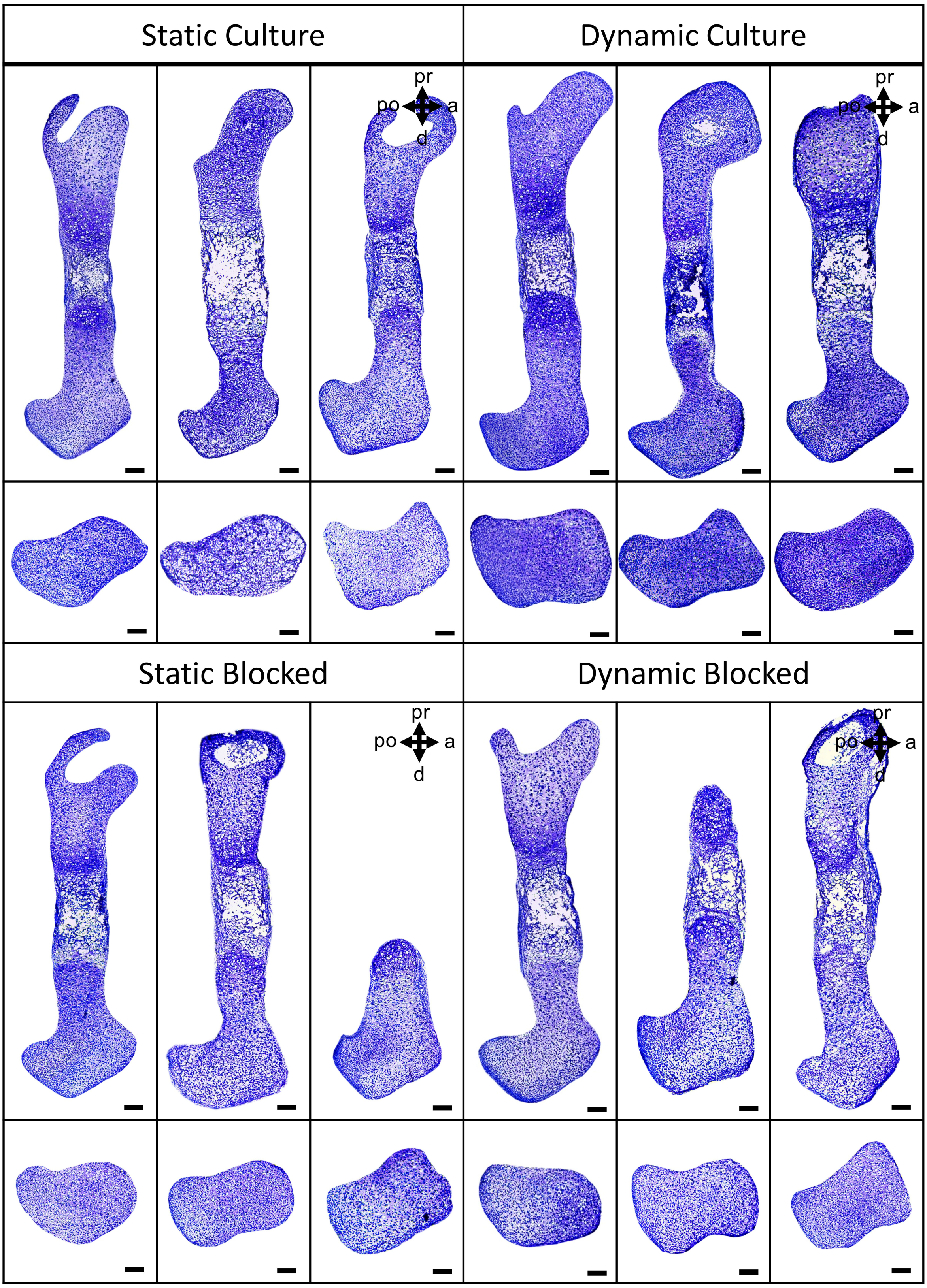


**S5: Glycosaminoglycans localization within all mouse embryo femora.** Each column represents contralateral limbs from a single embryo in two comparison groups. The static and dynamic vehicle control limbs (top row) are horizontally mirrored to aid comparison across the whole dataset. n=3 limbs per comparison group.

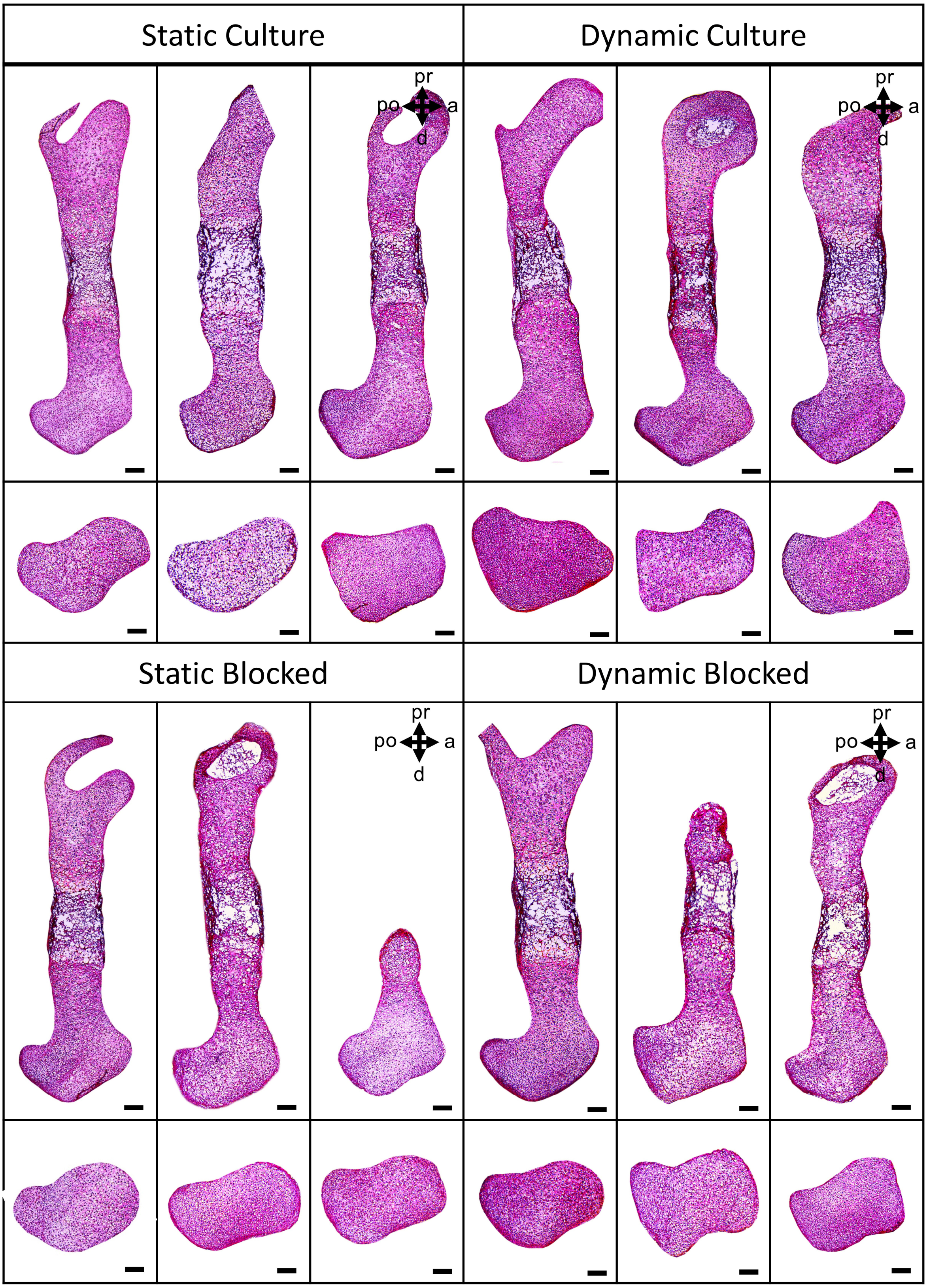


**S6: Collagen localization within all mouse embryo femora.** Each column represents contralateral limbs from a single embryo in two comparison groups. The static and dynamic vehicle control limbs (top row) are horizontally mirrored to aid comparison across the whole dataset. n=3 limbs per comparison group.

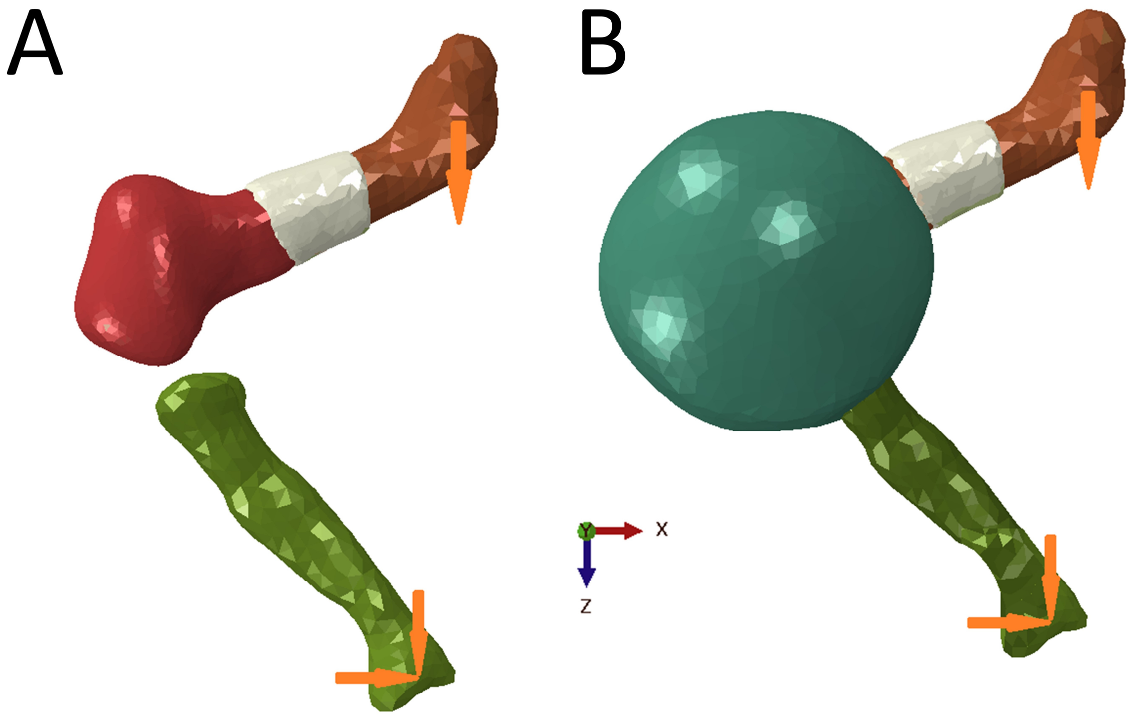


**S7: Finite element model set up.** (A) Principal Stress was calculated through Finite Element Analysis of an ideal limb (ABAQUS). Multiple OPT scans of limbs were collected, registered and separated into the distal femur and bone collar, with the proximal femur and tibia supplied by the most representative whole limb OPT scan. The relative orientation of the femur and tibia were adjusted to match the angle of limbs in the zero position from an image taken at day 6 of culture. (B) A solid spherical joint capsule was placed surrounding the joint space.
